# Historical shifts, geographic biases, and biological constraints shape mammal species discovery

**DOI:** 10.1101/2025.07.24.666531

**Authors:** Matheus de T. Moroti, Jhonny J. M. Guedes, Guilherme M. Missio, Giovana L. Diegues, Alexandra M. R. Bezerra, Mario R. Moura

**Affiliations:** Departamento de Biologia Animal, Instituto de Biologia, Universidade Estadual de Campinas, Campinas, SP, 13083-862, Brazil; Departamento de Ecologia, Instituto de Ciências Biológicas, Universidade Federal de Goiás – Campus Samambaia, Goiânia, GO, 74690-900, Brazil; Mastozoologia, Coordenação de Zoologia, Museu Paraense Emílio Goeldi, Belém, PA, 66077-830, Brazil; Departamento de Sistemática e Ecologia, Universidade Federal da Paraíba, João Pessoa, PB, 58051-900, Brazil

**Keywords:** Linnean shortfall, Mammalia, species descriptions, taxonomic practice, taxonomy

## Abstract

Species descriptions in taxonomy have become increasingly comprehensive, yet disparities persist across taxa and regions. We assess temporal trends in mammal species descriptions (1990–2025) using four proxies of comprehensiveness—counts of examined specimens and compared taxa, number of pages (only from the Methods/Results sections), and number of evidence lines (i.e., analytical tools and techniques). Using generalized linear models, we assessed how these proxies are explained by factors associated with species’ biology, geography, and taxonomic practice. Most new species derive from tropical regions, particularly rodents and bats, reflecting global discovery hotspots. Descriptions have grown more rigorous over time, with expanded specimen sampling, broader taxonomic comparisons, and integrative methods. However, disparities emerge along geographic and biological axes: descriptions from temperate regions incorporate more evidence lines, while small-bodied and tropical species (especially bats) remain understudied due to sampling biases and resource limitations. Body size inversely correlates with description length, as smaller species often require advanced diagnostics. Species-rich genera show greater comprehensiveness, likely due to heightened diagnostic scrutiny. Our findings highlight progress in taxonomic rigor but underscore persistent gaps tied to geography, body size, and accessibility of analytical tools. Addressing these disparities requires targeted investments in local capacity, equitable collaboration, and accessible methodologies to strengthen global taxonomic infrastructure and support conservation priorities.

## 1. Introduction

Species represent fundamental units in biodiversity science (Braby et al. 2024), where their names constitute testable hypotheses subject to revision in light of new data (Bremer et al. 1990; Raposo et al. 2017; Streicher et al. 2023; Gippoliti et al. 2024). Although taxonomic science is inherently dynamic and absolute stability is undesirable (Dominguez and Wheeler 1997; Dubois 2007), minimizing unnecessary changes in species identities can increase usability of biodiversity databases and improve scientific communication, data sharing, and conservation planning (Bertrand and Härlin 2006; Sullivan et al. 2023; Hamdan et al. 2024; Mahony et al. 2024). Following the development and popularization of integrative taxonomy, contemporary species descriptions tend to be more robust, incorporating more comprehensive diagnostic criteria (Sangster and Luksenburg 2015; Poulin and Presswell 2016; Guedes et al. 2024b) and contrasting with older descriptions, which generally relied on simpler diagnostics (Sangster and Luksenburg 2015; Poulin and Presswell 2016; Guedes et al. 2020). This helps explain the observed decline in synonymization rates over time (Solow et al. 1995; Alroy 2002; Baselga et al. 2008; Guedes et al. 2025), although this pattern may reflect a temporal bias (Alroy 2002) or changes in taxonomic practice (Lumpers vs. Splitters debate, Isaac et al. 2004). Understanding how temporal trends in species description comprehensiveness are related to improved sampling and analytical frameworks helps to inform best practices that can strengthen taxonomic science.

Comprehensive descriptions can help mitigate common pitfalls in taxonomic practices (Dayrat 2005; Wüster et al. 2021; Braby et al. 2024), such as oversplitting due to insufficient diagnostic criteria and failure to accurately distinguish natural variation within species (Isaac et al. 2004), or undersplitting resulting from overlooked phenotypic or genetic divergence (Chek et al. 2003; Hendrixson and Bond 2005; Mutanen et al. 2016). While the accumulated evidence highlights temporal shifts in taxonomic practices (Sangster and Luksenburg 2015; Padial and De la Riva 2021; Guedes et al. 2024b), recent studies also reveal pronounced spatial variation (De Queiroz 2007; Freeman and Pennell 2021; Padial and De la Riva 2021). A well-documented latitudinal taxonomic bias exists, wherein biodiversity-rich but resource-limited regions (e.g., the Tropics) often exhibit lower taxonomic resolution compared to temperate, high-income regions (Freeman and Pennell 2021; Diniz Filho et al. 2023; Guedes et al. 2025). This spatial variation likely emerges from differences in access to resources and tools, though international collaborations and open-access data initiatives have the potential to mitigate these imbalances (Nakamura et al. 2023; Carneiro et al. 2025; Weksler et al. 2025). Addressing these gaps is critical to ensuring robust taxonomic frameworks that support downstream research on fields dependent on taxonomic data (Diniz Filho et al. 2023; Lessa et al. 2024).

Because citation-based metrics are poor proxies of the comprehensiveness of taxonomic publications (Valdecasas et al. 2000; Valdecasas 2011; Pyke 2014), more direct metrics are needed to investigate trends in the robustness of species descriptions and to inform whether many taxonomic changes are likely to occur in the future across organismal groups. Using metrics like page counts and the number of evidence lines or number of techniques adopted (e.g., morphometrics, cytogenetics, internal anatomy, molecular analysis) in descriptions, studies have generally observed an increase in comprehensiveness of descriptions across birds (Sangster and Luksenburg 2015), reptiles (Guedes et al. 2024a), mollusks (Abreu et al. 2025), helminths (Poulin and Presswell 2016), and other taxa (Benton 2008; Pante et al. 2015). Here, we investigate how several proxies of comprehensiveness in species description have changed over time and assess their potential determinants.

We focus our assessment on recent mammals, a group with a relatively stable alpha taxonomy when compared to other vertebrates (e.g., amphibians and reptiles; see Moura and Jetz 2021; Guedes et al. 2024b), but yet still dynamic (Parsons et al. 2022). Specifically, we explore whether temporal variation is evident across four proxies of comprehensiveness in species descriptions, namely: (1) the number of specimens used to describe the new taxa, (2) count of taxa to which the new species is compared, (3) page counts, and (4) number of evidence lines used in the description. We examine whether variability in these proxies is related to species’ biology, geography, and taxonomic practice. Additionally, we inspect how accessibility to specific resources and tools is associated with the internationalization of species descriptions. Given that many mammalian taxa still lack comprehensive taxonomic studies (Feijó and Brandão 2022), and that most formal descriptions represent the splitting of one species into many (Isaac et al. 2004; Burgin et al. 2018), it is likely that several taxonomic changes lie ahead. Despite its focus on mammals, our study uses taxon-independent proxies for description comprehensiveness, which combined with its global scope, help yield broader implications for developing protocols and practices that can increase data quality and usage in biodiversity research (Mahony et al. 2024).

## 2. Material and methods

### 2.1. Data collection

We examined the descriptions of 1,116 mammal species (including terrestrial, freshwater, and marine) published between 1990 and 2025, covering 36 years of mammal discovery. Focusing on post-1990 descriptions minimize inconsistencies from historical variations in taxonomic standards, such as the cladistics and molecular systematics, ensuring comparable scientific and socioeconomic trends. We only considered “original” descriptions, disregarding redescriptions, revalidations, and other taxonomic works. Taxonomic nomenclature and species account follows the Mammal Diversity Database Version 2.1 (MDD 2025). For biogeographic regionalization, each species was assigned to a realm based on the coordinates of its type-locality, using the spatial data from Dinerstein et al. (2017).

For each species in our database, we extracted four proxies for the species description comprehensiveness: (1) number of specimens -type-specimens plus all referred specimens of the new species used to obtain diagnostic traits (type-series size and all referred specimens were highly correlated, Pearson’s *r* = 0.60, *p* < 0.001); (2) number of taxa compared - the count of close-related species directly compared with the new species; (3) number of pages - used as a proxy for the depth of taxonomic information. The presence of taxonomic keys, figures, or expanded results, for instance, typically represents a more detailed description and increases page and word count. We quantified this by dividing each page into quadrants and counting only those containing Methods and Results sections. For studies describing multiple species, the total count was divided by the number of species. Our fourth proxy was represented by the (4) number of evidence lines present in the formal description, scored from 1 to 9 lines of evidence based on the presence or absence of the following analyzed characters: (i) dentition (e.g. dental formula, tooth shape, etc), (ii) internal soft anatomy (e.g. internal morphology of the stomach, glans, etc.), (iii) external qualitative morphology (e.g. comparison of head and tail shapes), (iv) microscopic-based trichology, (v) coloration patterns, (vi) karyotype, (vii) external morphometry (e.g., head-body length, hind foot length, forearm length), (viii) osteological morphometry (from teeth, skull, mandible and other skeleton parts), and (ix) DNA (e.g. allozyme, mtDNA, nucDNA, multiLoci). These proxies showed low multicollinearity (Variance Inflation Factor, VIF < 1.5; Table S1) and limited pairwise correlations (Spearman’s *r* correlation range = 0.00-0.11; Fig. S1), indicating they capture relatively distinct aspects of description comprehensiveness.

To represent potential determinants of species description comprehensiveness, we used the seven variables: (1) year of species description; (2) number of authors; (3) average number of countries per author, the country counts derived from authors affiliations were divided by author counts to avoid sampling effects associated with multiauthored works (see Fig. S2 for pairwise correlation between authors and countries counts); (4) maximum body mass, obtained from original descriptions and from Moura et al. (2024), (5) per-genus species richness based on the year of species descriptions (i.e., for each species, we tallied the total number of currently accepted species described in the same genus until their year of description); (6) absolute latitude of type-locality; and whether description was based on (7) taxonomic revision, which was considered as such only if the work explicitly mentioned the term “revision” (or similar) in its title, abstract, keywords, or main text. To allow cross-group comparisons, we also recorded data on taxonomic ranks.

### 2.2. Statistical analyses

Temporal trends in our proxies of the species description comprehensiveness (number of lines of evidence, number of pages, number of specimens, and number of taxa compared) were investigated using Spearman’s (*r_s_*) correlation between the year of publication and annual means for the metrics analyzed across species described in each year. We assessed multicollinearity among continuous predictors using the Variation Inflation Factors (VIF; Mansfield and Helms 1982), where strong multicollinearity is often attributed to variables holding VIF > 10 (Kutner et al. 2005). All variables in our analysis had VIF values below 2 (Table S2), so none were excluded.

We assessed potential determinants of species description comprehensiveness, using generalized linear models (GLMs; Zuur et al. 2009). We used a negative binomial error distribution to analyze the number of specimens and number of taxa compared, both of which are count data. The number of pages was calculated as a continuous variable, so we applied a log_10_-transformation and modelled it using a GLM with a Gaussian error distribution. Although the number of evidence can be considered a count variable, given its hump-shaped distribution (Fig. S1), we used both negative binomial and Gaussian distributions, comparing model fit and selecting the one with the lowest AIC values (Burnham and Anderson 2002). Each response variable was modelled against year of species description, number of authors (log_10_-transformed), average number of countries per author (log_10_-transformed), body mass (log_10_-transformed), per-genus species richness (log_10_-transformed), and occurrence of taxonomic revision. Continuous predictors were *z*-transformed before modelling, allowing comparisons of their effect sizes. Because of varying amounts of missing data among response and predictor variables, GLMs were performed with datasets differing slightly in the number of modelled species: number of lines of evidence (*n* = 937 species); number of pages (*n* = 950), number of specimens (*n* = 933); and number of taxa compared (*n* = 942). All analyses were performed in the R software version 4.3.1 (R Core Team 2023).

All models were fitted using the package *MASS* (Venables and Ripley 2002). We assessed explained variation through Nagelkerke’s pseudo-R^2^ for negative binomial models (response variables ‘number of taxa compared’ and ‘number of specimens’) and the coefficient of determination (R^2^) for the Gaussian model (response variables ‘number of evidence’ and ‘number of pages’), both computed with the R package *performance* (Lüdecke et al. 2021). We assessed the necessity of phylogenetic regression models by investigating the phylogenetic autocorrelation of model residuals through Moran’s *I* correlograms, computed across 14 distance classes (Revell 2010). Correlograms were derived from the average results of 100 fully sampled phylogenies for mammals (Upham et al. 2019), with trees trimmed to include only modelled species. Phylogenetic autocorrelation analyses were conducted with the R packages *phylobase* (Bolker et al. 2020) and *phylosignal* (Keck et al. 2016).

To further investigate if accessibility of costly analytical tools is reflected across international collaboration, we applied a Kruskal-Wallis analysis to test whether the average number of countries per authors differed among descriptions using (i) molecular biology, (ii) taxonomic review, or (iii) other evidence lines. *Post-hoc* comparisons were conducted using Dunn’s test, with pairwise contrasts adjusted for multiple testing using the Bonferroni correction method (Bonferroni 1936). All analyses were conducted using the *rstatix* R package (Kassambara 2023). See Data Availability section for raw data and R code (Moroti et al. 2025).

## 3. Results

Most mammal species described between 1990 and 2025 were from Africa, South America, Southeast Asia, and islands near Australia (Fig. 1). The Afrotropical realm stands out in terms of primate descriptions, whereas the few descriptions from the Nearctic were mostly from rodents and Eulipotyphla (e.g., hedgehogs, moles, true shrews). The majority of the 1,116 species described comprised rodents (41%, n = 458) and bats (26.2%, n = 293), a trend observed globally as well as across most biogeographic realms. Descriptive statistics for comprehensiveness metrics across all mammals, rodents, bats, and when considering mammals excluding bats and rodents, are summarized in Table 1.

**Figure 1.**
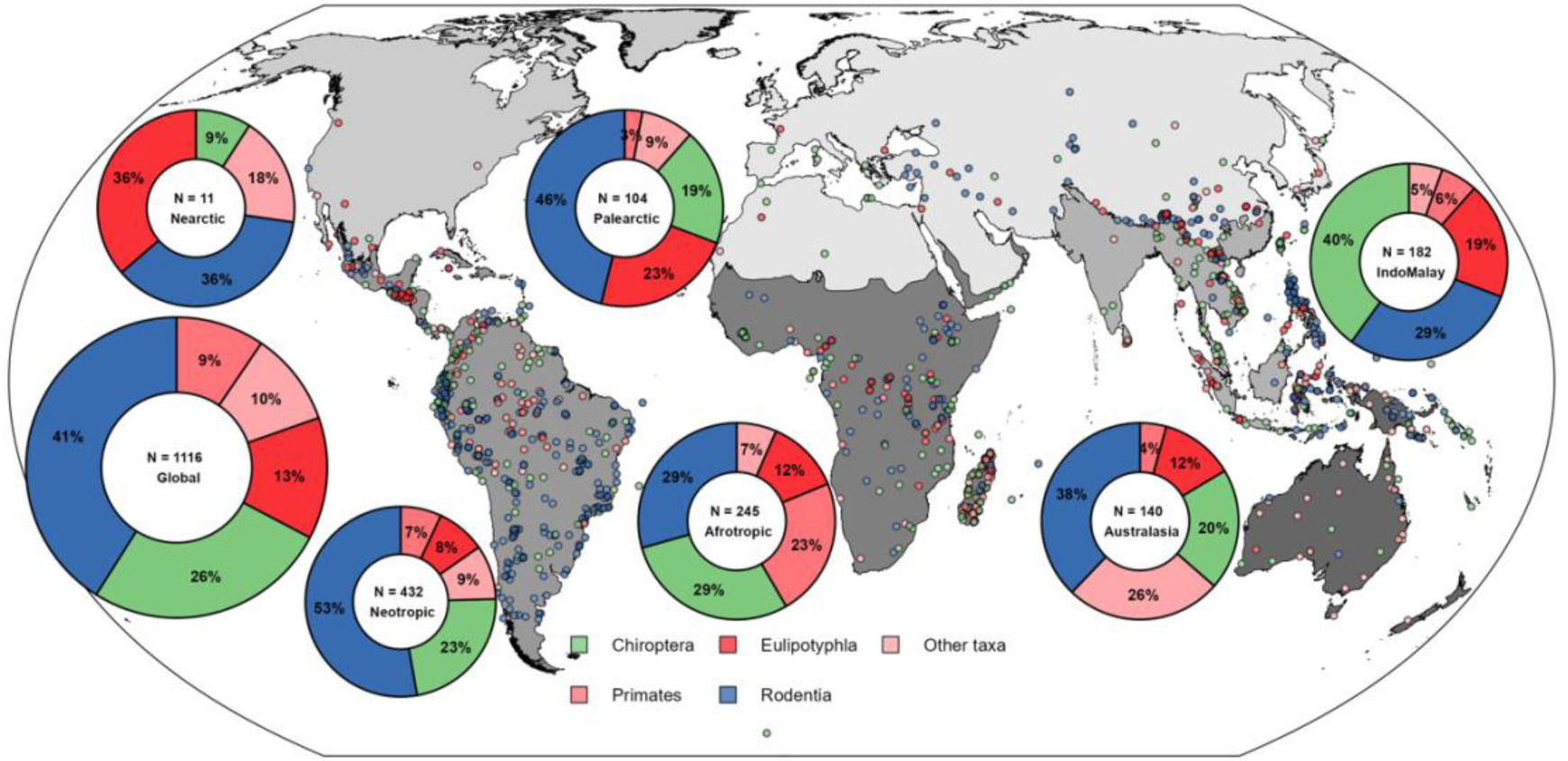
Global distribution of mammal species described between 1990 and 2025. Each point on the map represents a species’ type locality, while donut charts indicate the proportion of species described within each taxonomic order. Points are coloured according to their respective taxonomic order, with rodents in blue, bats in green, and the different shades of red representing other mammals. To improve visualization, taxonomic orders holding less than 5% of species descriptions were grouped under ‘Other taxa’ label in the donut charts. The total sample size (N) is lower than the 1,116 species described during this period due to missing type-locality data for some species. Two species occurring in the Oceania realm were suppressed due to low representation.

**Table 1.** Descriptive statistics for metrics of species descriptions’ comprehensiveness. For each group, we report the median and the range (minimum–maximum) for the number of evidence lines, number of specimens, number of pages, and number of taxa compared.

|  | All mammals | Non-bats &<br>non-rodents | Bats | Rodents |
| --- | --- | --- | --- | --- |
| N. of evidence lines | 5 (1–8) | 5 (1–7) | 5 (1–8) | 5 (1–8) |
| N. of specimens | 8 (1–409) | 7 (1–345) | 7 (1–327) | 9 (1–409) |
| N. of pages | 7.5 (0.23–179) | 6.5 (0.23–48.2) | 8.1 (0.8–179) | 8.1 (1–130) |
| N. of taxa compared | 5 (0–77) | 4 (0–46) | 5 (1–34) | 4 (0–77) |

### 3.1. Temporal trends in mammal description comprehensiveness

The annual mean number of evidence lines used in species descriptions increased when analyzing all mammals (*r_s_* = 0.75; *p* < 0.001), non-bat & non-rodent mammals (*r_s_* = 0.51; *p* = 0.006), bats (*r_s_* = 0.63; p < 0.001) and rodents (*r_s_* = 0.64; *p* < 0.001; Fig 2a). There was no change in the number of pages over time (Fig. 2b), but an increase in the number of specimens when considering all mammals together (*r_s_* = 0.49; *p* = 0.02, Fig. 2c) and non-bat & non-rodent mammals (*r_s_* = 0.44; *p* = 0.03). The number of taxa compared also increased for all mammals combined (*r_s_* = 0.61; *p* < 0.001, Fig. 2d), non-bat & non-rodent mammals (*r_s_* = 0.56; *p* < 0.001) and for bats separately (*r_s_* = 0.7; *p* < 0.001), but not for rodents (*r_s_* = 0.25; *p* = 0.71). In addition, the incorporation of molecular analyses in species descriptions has grown over time, with a noticeable increase in the proportion of taxa described annually using molecular data (Figs S3-S4).

**Figure 2.**
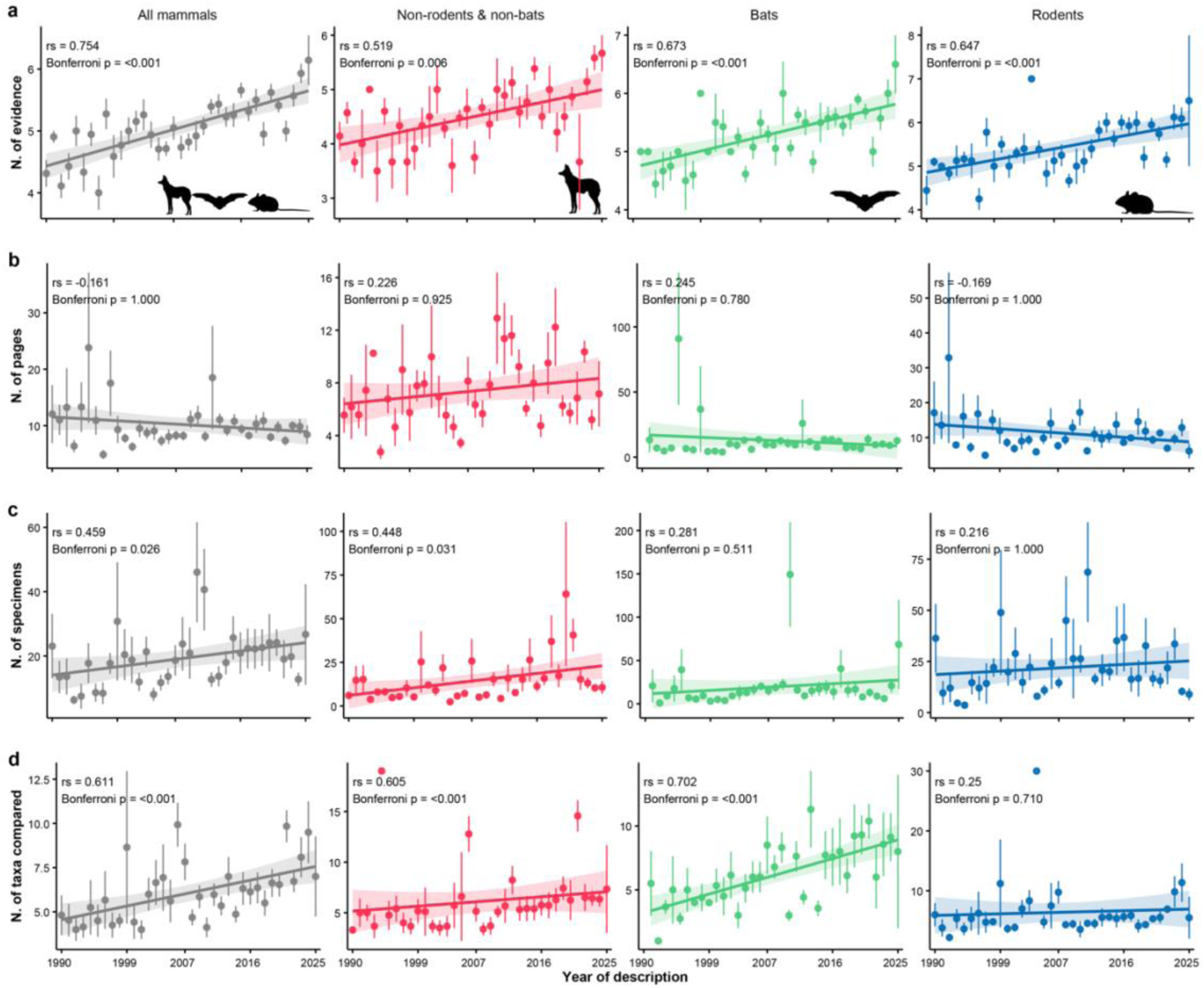
Variation in proxies of mammal species descriptions comprehensiveness over time (1990–2025). Panels show overall trends for all mammals (gray), non-rodents and non-bats (red), bats (green), and rodents (blue). Proxies include the (a) number of evidence used in species descriptions, (b) number of pages in taxonomic description papers, (c) number of specimens examined in species descriptions, and (d) number of taxa compared to distinguish new species. rs denotes the Spearman’s correlation between the year of description and annual means for each variable. We applied Bonferroni corrections to p-values to account for multiple comparisons. Vertical lines represent standard errors, and shaded areas around the regression lines represent 95% confidence intervals.

### 3.2. Internationalization of mammal descriptions

We found a temporal decrease in the average number of countries per author for all taxonomic groups (Fig. 3). In addition, this ‘internationalization indicator’ differed across descriptions using analytical tools, with studies using molecular biology and taxonomic review showing lower values of average number of countries per author. In bats and rodents, only molecular-based descriptions involved fewer countries relative to other lines of evidence.

**Figure 3.**
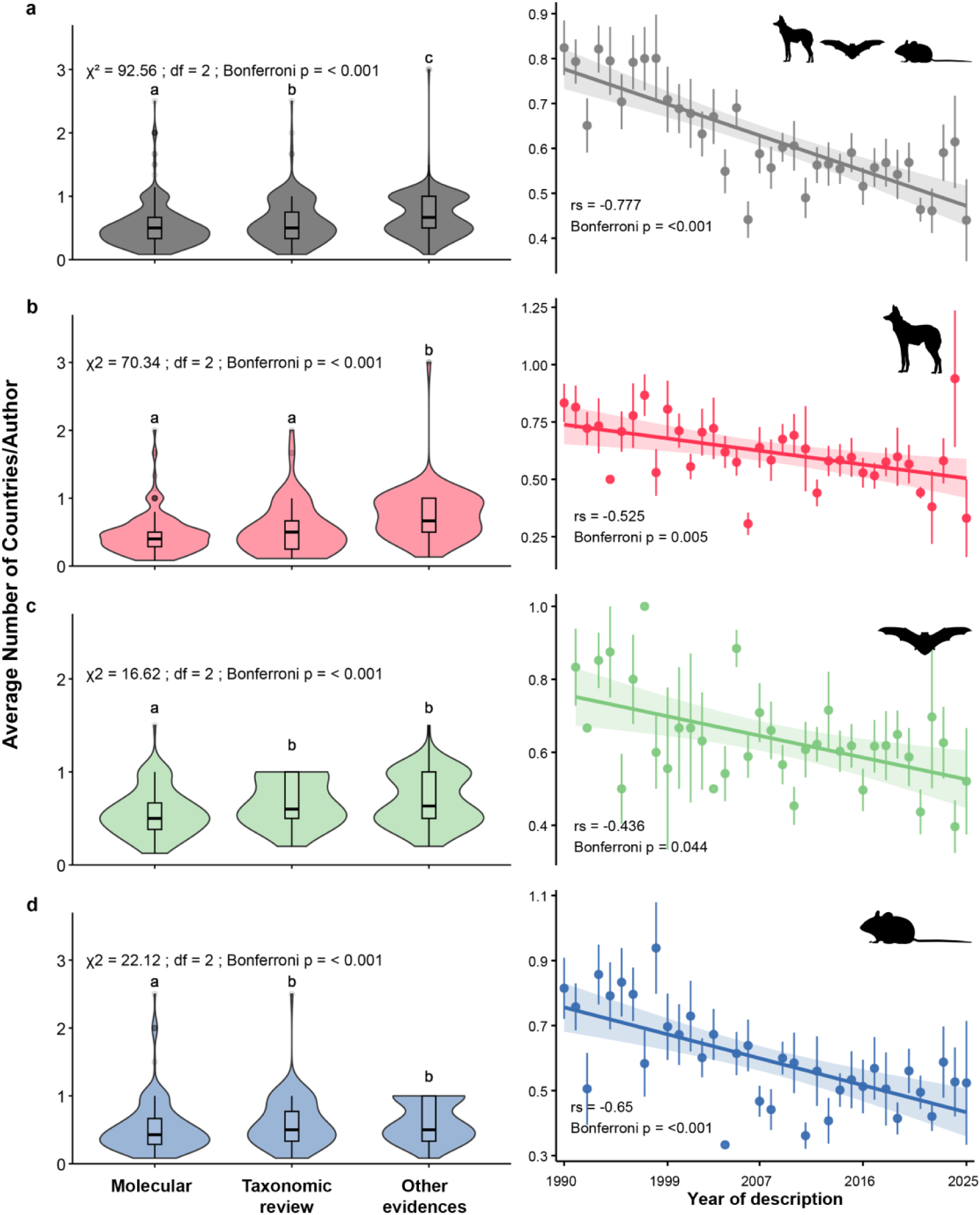
Average number of countries per author involved in mammal species descriptions across different lines of evidence and over time. Patterns are shown for (a) all mammals, (b) non-bats and non-rodents, (c) bats, and (d) rodents. ‘Other evidences’ refers to descriptions that do not include molecular diagnoses and were not derived from taxonomic revisions. Molecular approaches exhibit different levels of internationalization compared to taxonomic reviews and other evidence types across mammals, bats, and rodents. In the violin plot, post-hoc comparisons were performed using estimated marginal means, with pairwise contrasts adjusted for multiple testing using the Bonferroni method. Groups that do not share letters are significantly different from each other.

The use of molecular data in descriptions varies across mammalian orders, with Pilosa (4 spp.), Carnivora (5 spp.), Artiodactyla (19 spp.), Peramelemorphia (5 spp.), Macroscelidea (3 spp.), Didelphimorphia (25 spp.), Rodentia (n = 458 spp.), and Eulipotyphla (146 spp.) frequently incorporating genetic evidence (more than 50% described species). In comparison, Dasyuromorphia (13 spp.) lacked such data entirely. International collaborations were common in molecular-based descriptions, except in Lagomorpha, with nine descriptions associated with a single country (Fig. 4a). Taxonomic revisions contributed substantially to new species descriptions in Pilosa, Monotremata, and Carnivora (> 50%), but were rare in species-rich groups like Rodentia and Chiroptera and absent in depauperate orders (e.g., Peramelemorphia, Paucituberculata, Microbiotheria, Macroscelidea, Lagomorpha, and Hyracoidea). While international collaboration in revision-based descriptions was idiosyncratic, primates and afrosoricids exhibited broader multinational involvement—i.e., >75% of descriptions involving two or more countries (Fig. 4b).

**Figure 4.**
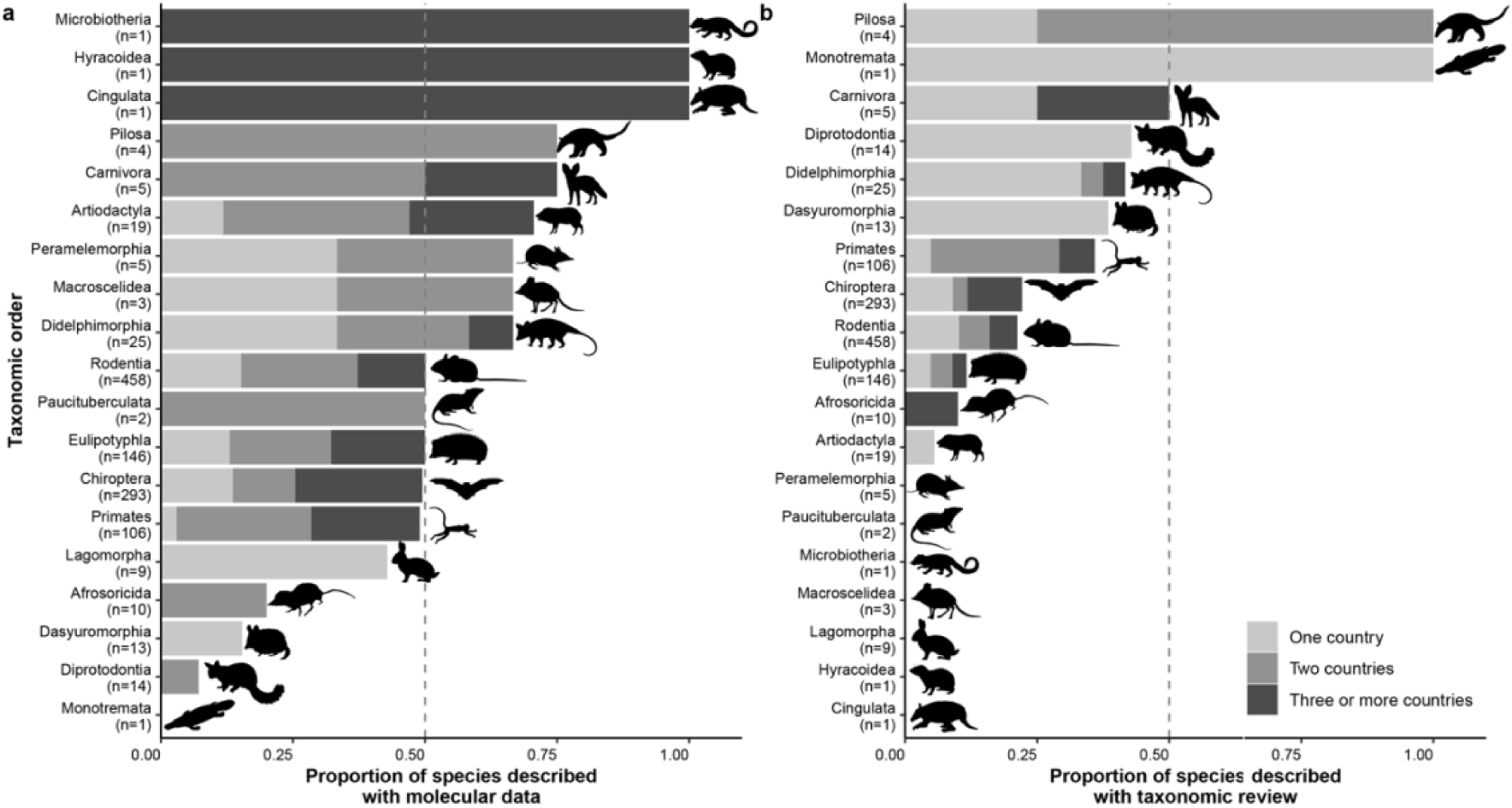
Proportion of species descriptions using (a) molecular analysis or (b) taxonomic review for each mammal order from 1990–2025. The bars indicate the proportion of the total number of species described per order over this period. The different shades of gray indicate the proportion of these descriptions involving one, two, and three or more countries.

### 3.3. Determinants of comprehensiveness in mammal descriptions

GLMs explained 2–30% of the variation in taxonomic comprehensiveness metrics (Fig. 5). Across all four proxies tested, the number of taxa compared emerged as the best-explained one for descriptions in all mammals and rodents, whereas the number of specimens was the most explained metric in non-bat & non-rodent mammals and bats. We did not find phylogenetic structure in GLM residuals, as indicated by phylogenetic correlograms (Fig. S5).

**Figure 5.**
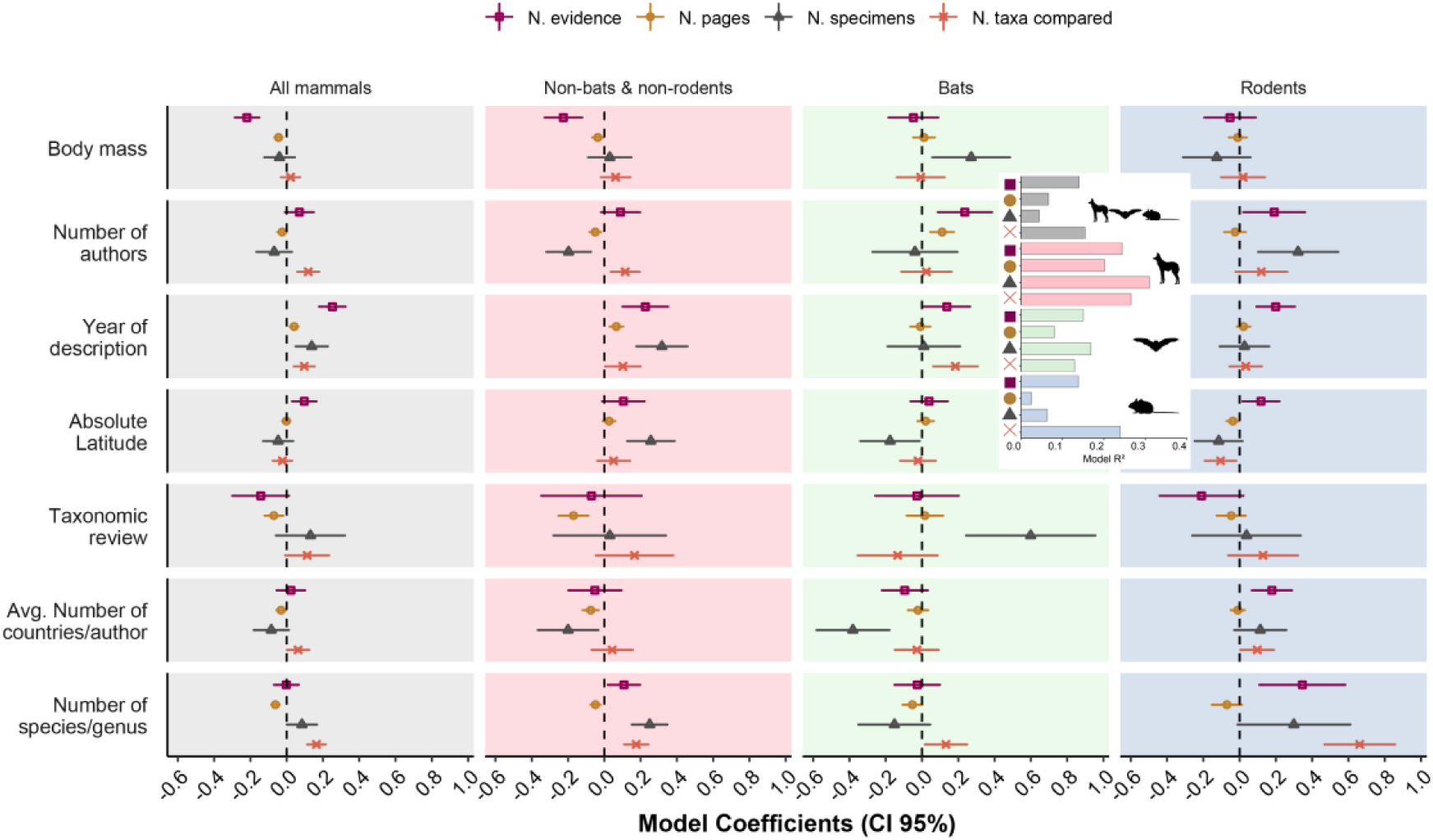
Influence of putative drivers on proxies of mammal species description comprehensiveness. Standardized coefficients from the generalized linear models fit to all species and for bats and rodents separately are shown. Horizontal lines denote the 95% confidence intervals around each coefficient. Predictors with significant effect sizes (p-value < 0.05) are those whose horizontal lines do not cross 0. Explained variation is presented (Model R²) as Nagelkerke’s pseudo-R² for negative binomial models (Number of taxa compared and Number of specimens) and as the coefficient of determination (R²) for the Gaussian model (Number of evidence and Number of pages).

We found the year of description, number of species/genus, and absolute latitude among the most important predictors of species description comprehensiveness. Recent descriptions used more evidence, referred specimens, compared taxa, and (to some extent) more pages across all taxonomic mammalian orders and non-bat & non-rodent mammals, although such effect was less pronounced in bats (Fig. 5). Species-rich genera correlated with more specimens and compared taxa overall, except for bats specimens. Our results revealed a latitudinal gradient, with the number of evidence lines decreasing in mammal descriptions from tropical regions. We also observed fewer specimens in descriptions of non-rodent and non-bat species towards the Tropics. However, more taxa are compared in the Tropics for rodents.

Other predictors affected only some taxonomic groups. For instance, we found fewer evidence lines and pages in descriptions of small-sized mammals that are neither rodents nor bats, and all mammals when considering them together. In bats, fewer specimens were used to describe small-sized species, but descriptions produced via taxonomic review generally included more specimens. For non-bat and non-rodent mammals, taxonomic reviews tended to promote descriptions with fewer pages.

The two remaining predictors, number of authors and average number of countries per author (an indicator of internationalization), showed more contrasting trends across taxonomic groups and proxies of description comprehensiveness. Non-rodent and non-bat species had fewer specimens in multi-authored descriptions. In contrast, bat descriptions with more authors included more evidence, and international collaborations had limited influence on description comprehensiveness, as descriptions of non-rodent and non-bat mammals with multinational authorship consistently included fewer specimens.

## 4. Discussion

Most new mammal species described between 1990 and 2025 come from Tropical regions, particularly rodents and bats. This pattern aligns with mammal discovery hotspots (Moura and Jetz 2021; Parsons et al. 2022) and corresponds to the highest diversity of both orders in these regions (Abreu et al. 2024; MDD 2025). Although the rate of new mammal descriptions remains slower than in ectothermic tetrapods (Moura et al. 2018), hundreds of species await formal descriptions (Moura and Jetz 2021; Parsons et al. 2022), with 68.8% increase in recognized species since the 1980s (∼65 new species/year; Burgin et al. 2025). Many additions result from taxonomic splits, reflecting taxonomic uncertainties often rooted in species descriptions of limited comprehensiveness, likely due to the historical unavailability or accessibility of tools that are now commonly used in taxonomic research (Sangster and Luksenburg 2015; Poulin and Presswell 2016; Guedes et al. 2024b). We find a clear temporal progression toward more rigorous practices in mammal descriptions, including an increased use of multiple evidence lines, expanded sampling of referred specimens, and broader taxonomic comparisons (Fig. 2). Our results reveal substantial heterogeneity in determinants of species description comprehensiveness, likely associated with taxon-specific sampling challenges.

The increase over time in comprehensiveness of mammal species descriptions has mirrored trends observed across various taxa (Sangster and Luksenburg 2015; Poulin and Presswell 2016; Guedes et al. 2024b). While this pattern aligns with the principles of integrative taxonomy (Dayrat 2005; Will et al. 2005), it also shows that many taxonomists were already applying integrative approaches before the formalization of the concept (Valdecasas et al. 2008). The higher comprehensiveness of recent descriptions can be attributed to technological advancements and reduced costs, which have democratized access to analytical tools (Nakamura et al. 2023; Guedes et al. 2024b), like micro-CT scanning, DNA sequencing, and automated platforms for taxonomic analysis (Thomer et al. 2018; Grenié et al. 2023; e.g., Blackburn et al. 2024). Taken together, these technological advances have accelerated species descriptions and expanded the scope and depth of taxonomic knowledge, contributing to a more comprehensive understanding of biodiversity. Despite improvements in access to analytical tools and resources, disparities still persist (e.g., Abreu et al. 2025), as evidenced by a greater number of evidence lines in mammal descriptions from temperate regions (Fig. 5), where wealthy economies predominate relative to low latitudes.

While descriptions of new mammal species now involve more authors and countries overall (Fig. S6), the growth in authorship size has outpaced international collaboration (Fig. 3). Consequently, we find a lower average number of countries per author in recent descriptions, which is also noticeable across molecular-based descriptions. These findings may relate to novel standards in authorship criteria of recent descriptions (Allen et al. 2019; Lin 2024). For instance, more collectors of mammal type-specimens have authored recent descriptions (Fig. S7). This suggests that recent species descriptions may be based on newly collected specimens obtained through projects led by these collectors, or that there is an increasing recognition of the contributions made by fieldwork personnel. In addition, the somewhat limited internationalization of molecular-based descriptions corroborates DNA sequencing in taxonomic research, which is nowadays more affordable by researchers in biodiversity-rich regions in the Global South. For instance, the Human Genome Project (1990-2003) required 3 US$ billions, whereas advances like next−generation sequencing reduced costs to less than $1,000 per genome (Schmidt and Hildebrandt 2017; Wetterstrand 2025). Although this shift is evident in the rising prevalence of molecular-based diagnoses in mammal descriptions (Figs S4-S5), the advent of next-generation sequencing has introduced greater methodological heterogeneity and costly approaches that are not necessarily accessible to Global South researchers.

Among the most consistent findings of this study is the higher comprehensiveness of descriptions made within speciose genera. On the one hand, this finding may reflect a sampling effect, as species-rich genera necessitate more extensive within-group comparisons as additional species accumulate over time (Fig. 2). On the other hand, alpha taxonomy is inherently comparative, and distinguishing closely related species becomes particularly challenging in species-rich genera (Gaston et al. 1995), where greater phenotypic and genetic variation may require more specimens to accurately delineate species. Indeed, we found that more speciose genera from all groups analyzed together, or separately, involve more taxa compared. Similarly, more evidence is used for diagnosis in rodents and non-bats & non-rodents. On the other hand, only in mammals analyzed together, and non-bats & non-rodents have more specimens included. This pattern may be related to the difficulty of collecting specimens from small and nocturnal, difficult-to-sample animals, such as rodents and bats (Meiri 2018; Moura and Jetz 2021).

Species descriptions of larger mammals contain fewer evidence lines and are shorter than those of smaller species, especially in non-flying mammals. While larger species’ morphologies are typically diagnosable without advanced imaging, smaller species often require it for accurate identification (Ziegler and Sagorny 2021). Thus, body size shows a negative relationship with evidence lines, contrasting to its positive associations with species description probability and data availability (Meiri 2018; Moura and Jetz 2021). The inverse association between body mass and description length appears to be mediated through the number of evidence lines. That is, small species demand more pages to document their additional evidence lines, a pattern supported by the moderate but significant correlation between page count and evidence lines (Fig. S1). Notably, body mass influenced specimen availability only in bats, once again reflecting taxon-specific sampling challenges. Unlike bats, which face size-biased mist-netting limitations, non-flying mammals (especially rodents) are more efficiently collected through passive methods like pitfalls and baited live traps (Tobler et al. 2008), with minimal size-related bias (Williams and Braun 1983; Bovendorp et al. 2017). These sampling limitations result in fewer specimens of bats relative to non-flying mammals, as evidenced by GBIF’s global holdings of preserved specimens (1.16 million bats versus 3.6 million rodents and 1.89 million other non-flying mammals; GBIF 2025), although these figures may also reflect the global distribution of known species by order (MDD 2025).

Our study highlights the significant progress in the comprehensiveness of mammal species descriptions since the 1990s, driven by technological advancements and the broader adoption of integrative taxonomic practices. Additionally, our findings confirm that taxonomic practices align with the well-documented latitudinal gaps in taxonomic resolution (Freeman and Pennell 2021; Guedes et al. 2025), with implications that extend beyond mammalogy. The core question of how taxonomic effort shapes our understanding of biodiversity is universal. The pattern we uncover in mammals, which are driven by socioeconomic inequalities, taxon-specific sampling biases, and the broader taxonomic impediment (i.e., systemic barriers limiting both taxonomic output and end-user reliability; Ebach et al. 2011; Moura et al. 2025), exemplifies the systemic issues faced in biodiversity systematics. Bats, rodents, and potentially other small-sized Tropical organisms face compounded challenges, from methodological and sampling constraints to underrepresentation in either molecular diagnoses or revisionary studies (Guedes et al. 2024a; Abreu et al. 2025). Addressing these challenges requires targeted investments in building local capacity and inclusive collaboration frameworks (Nakamura et al. 2023; Carneiro et al. 2025; Moura et al. 2025). In the absence of such interventions, the interoperability of taxonomic data risks being delayed (Sandall et al. 2023), ultimately undermining research on conservation, systematics, and evolution.

## Acknowledgments

We thank Pedro C. Estrela for valuable comments on a previous version of our manuscript. JJMG thanks Conselho Nacional de Desenvolvimento Científico e Tecnológico (CNPq, proc. 381394/2024-7) and Coordenação de Aperfeiçoamento de Pessoal de Nível Superior (CAPES, proc. 88887.478942/2020-00) for research grants. We also acknowledge support from Rio de Janeiro Foundation (FAPERJ) for grant to ARB (E-26/203.454/2023), and São Paulo Research Foundation (FAPESP) for grants to MTM (#2023/14506-5), GMM (#2023/16169-6), GLD (#2022/14674-2), and MRM (#2021/11840-6 and #2022/12231-6). This work is also a contribution of the National Institute of Science and Technology (INCT) in Ecology, Evolution, and Biodiversity Conservation funded by CNPq (grant 465610/2014-5/ 409197/2024-6).

## Data Sharing

The R-script and raw dataset used in the analyses of this study is available at Zenodo Digital Repository (Moroti et al. 2025) as well as on GitHub (https://github.com/jhonny-guedes/mammal_desc_trends)

