## Supplementary material for "Historical shifts, geographic biases, and biological constraints shape mammal species discovery"

**Table S1. Variance Inflation Factor (VIF) values of predictors for the proxies of species description comprehensiveness.** All variables in our analysis had VIF values below 1.5, indicating they capture relatively distinct aspects of description comprehensiveness.

| **Proxies** | **All mammals** | **Non-bats & non-rodents** | **Bats** | **Rodents** |
| --- | --- | --- | --- | --- |
| Number of specimens | 1.006 | 1.025 | 1.018 | 1.001 |
| Number of taxa compared | 1.004 | 1.024 | 1.006 | 1.005 |
| Number of authors | 1.016 | 1.069 | 1.010 | 1.009 |
| Number of evidence | 1.024 | 1.099 | 1.021 | 1.009 |

**Table S2. Variance Inflation Factor (VIF) values of predictors for the proxies of species description comprehensiveness.** All variables in our analysis had VIF values below 2, indicating they capture relatively distinct aspects of predictors in description comprehensiveness.

| **Predictors** | **All mammals** | **Non-bats & non-rodents** | **Bats** | **Rodents** |
| --- | --- | --- | --- | --- |
| Year of description | 1.240 | 1.288 | 1.325 | 1.331 |
| Body mass | 1.109 | 1.385 | 1.047 | 1.005 |
| Number of authors | 1.516 | 1.674 | 1.529 | 1.815 |
| Avg. N. of countries per author | 1.402 | 1.569 | 1.248 | 1.543 |
| Genus richness | 1.084 | 1.320 | 1.077 | 1.066 |
| Absolute latitude | 1.061 | 1.165 | 1.067 | 1.060 |

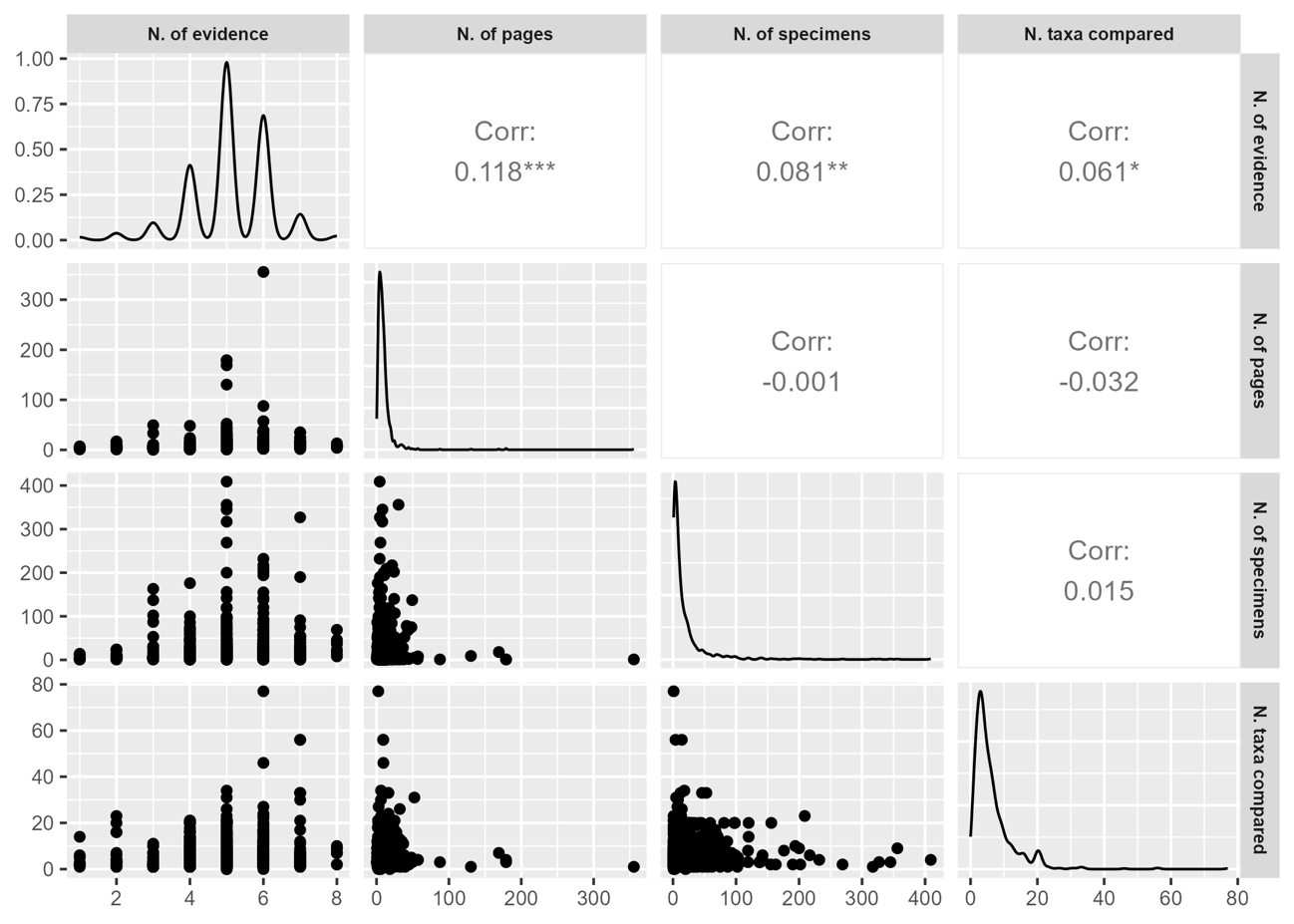

**Figure S1. Pairwise correlations among taxonomic robustness metrics. Exploratory analysis of pairwise correlations (Spearman's) between quality metrics (Number of evidence, number of pages, number of specimens, and number of taxa compared).** All correlations were weak (*r_s_* < 0.4), indicating that these metrics vary relatively independently in descriptions of mammalian species.

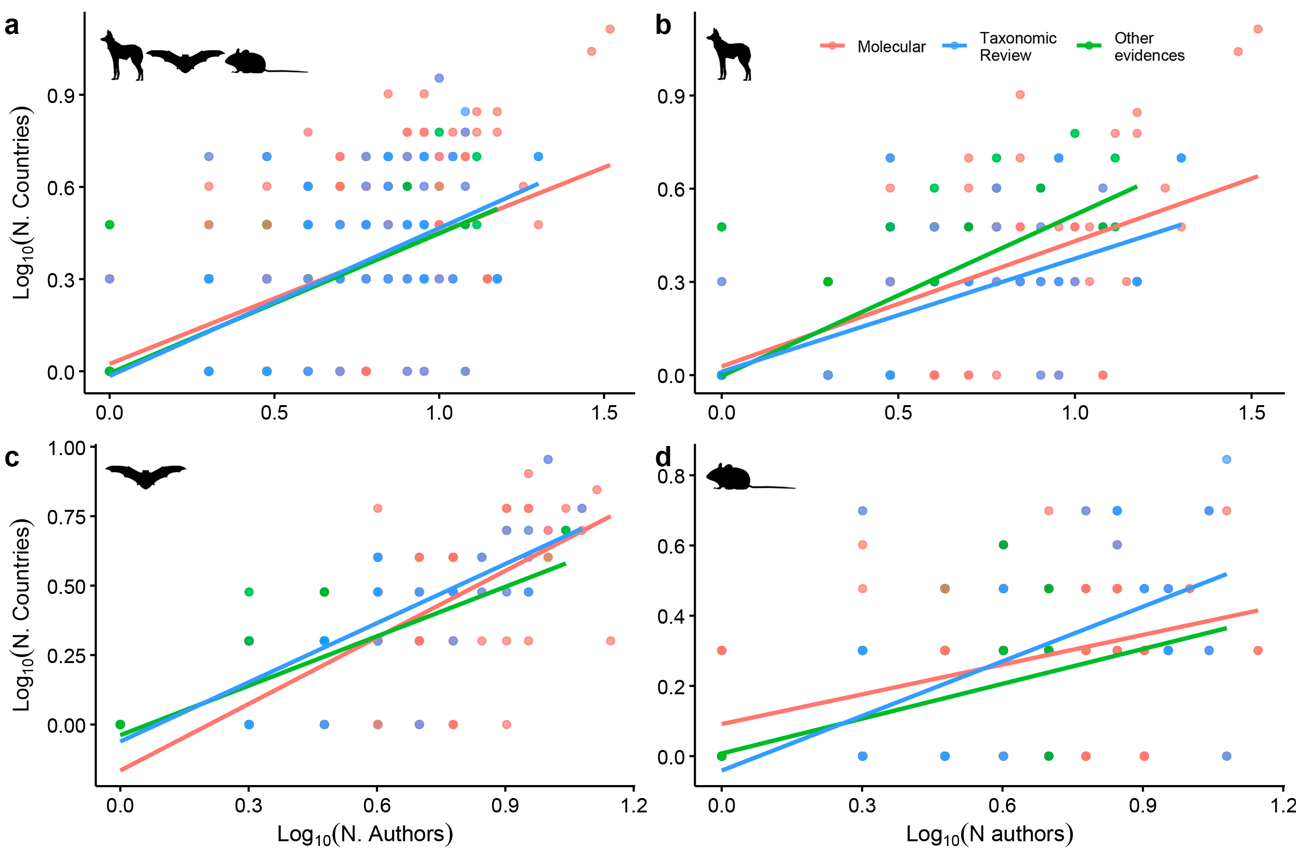

**Figure S2.** **Relationship between the number of authors (x-axis Log10 transformed) and the number of countries involved in different species description approaches (y-axis Log10 transformed).** Panels show (a) all mammal species; (b) non-bats & non-rodents; (c) bats; and (d) rodents. Across all groups, there is a positive trend in the relationship between the two variables.

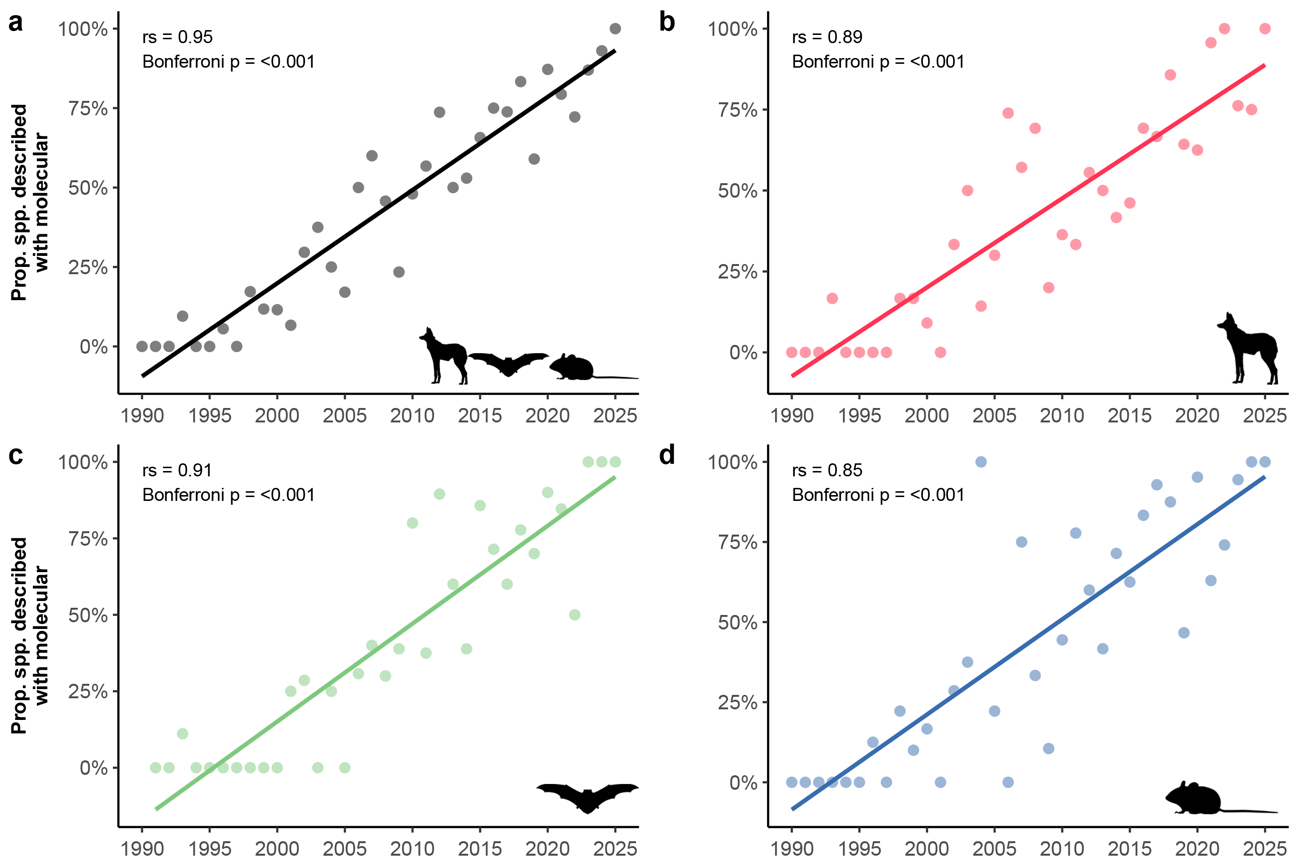

**Figure S3.** **Temporal trends in molecular analysis in mammal species descriptions from 1990–2025.** Panels show (a) all mammal species; (b) non-bats & non-rodents; (c) bats; and (d) rodents. Across all groups, there was an increase over time in the proportion of species described using molecular methods.

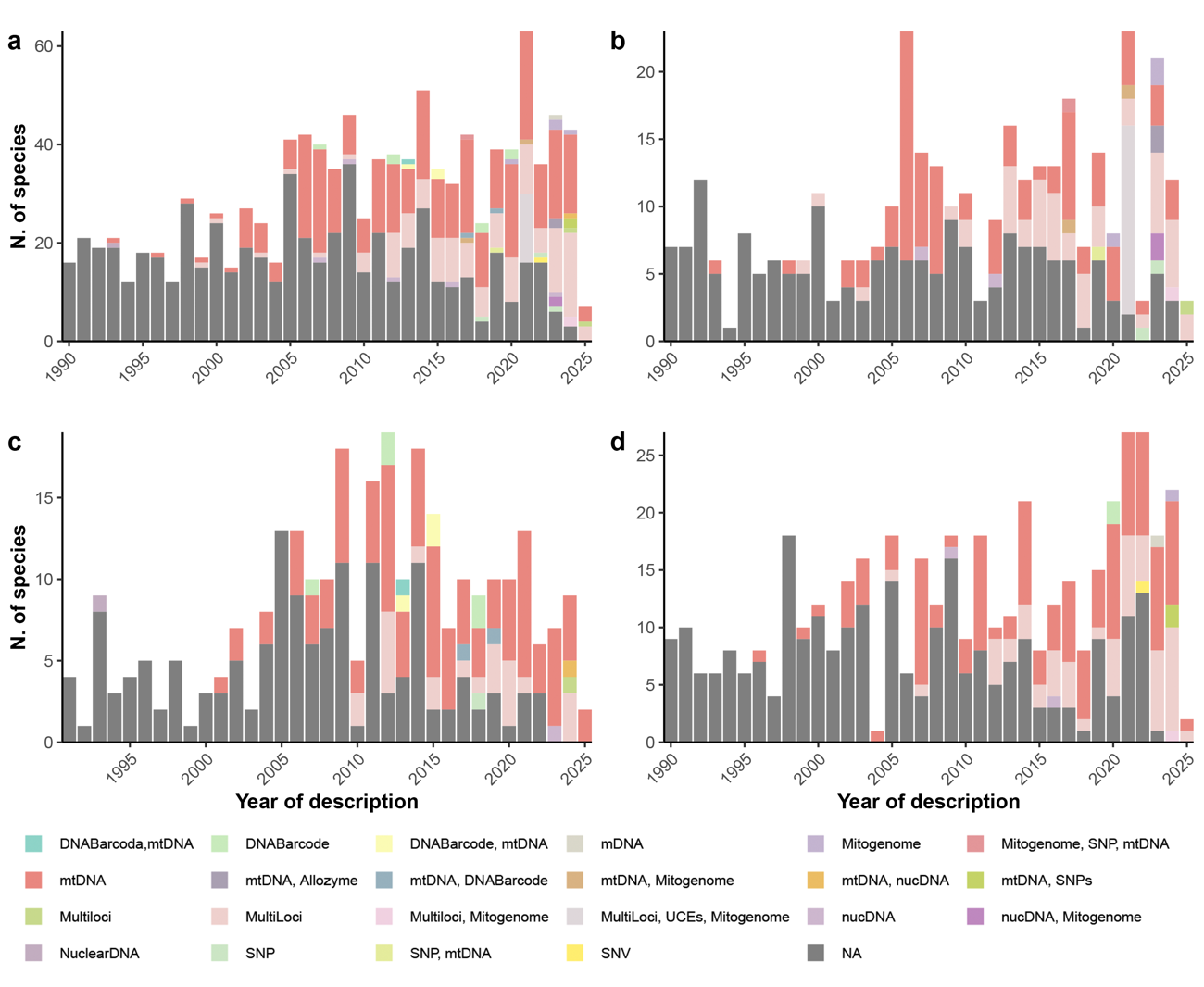

**Figure S4.** **Molecular methods used over time for (a) all mammal species; (b) non-bats & non-rodents; (c) bats; and (d) rodents.** In molecular methods, SNP stands for single-nucleotide polymorphism; UCE stands for ultraconserved element; and NA means no molecular data was used. The mtDNA and multilocus approaches were generally the predominant methods, with few descriptions based on allozymes, DNA barcode, mitogenome, SNPs, or nuclear DNA.

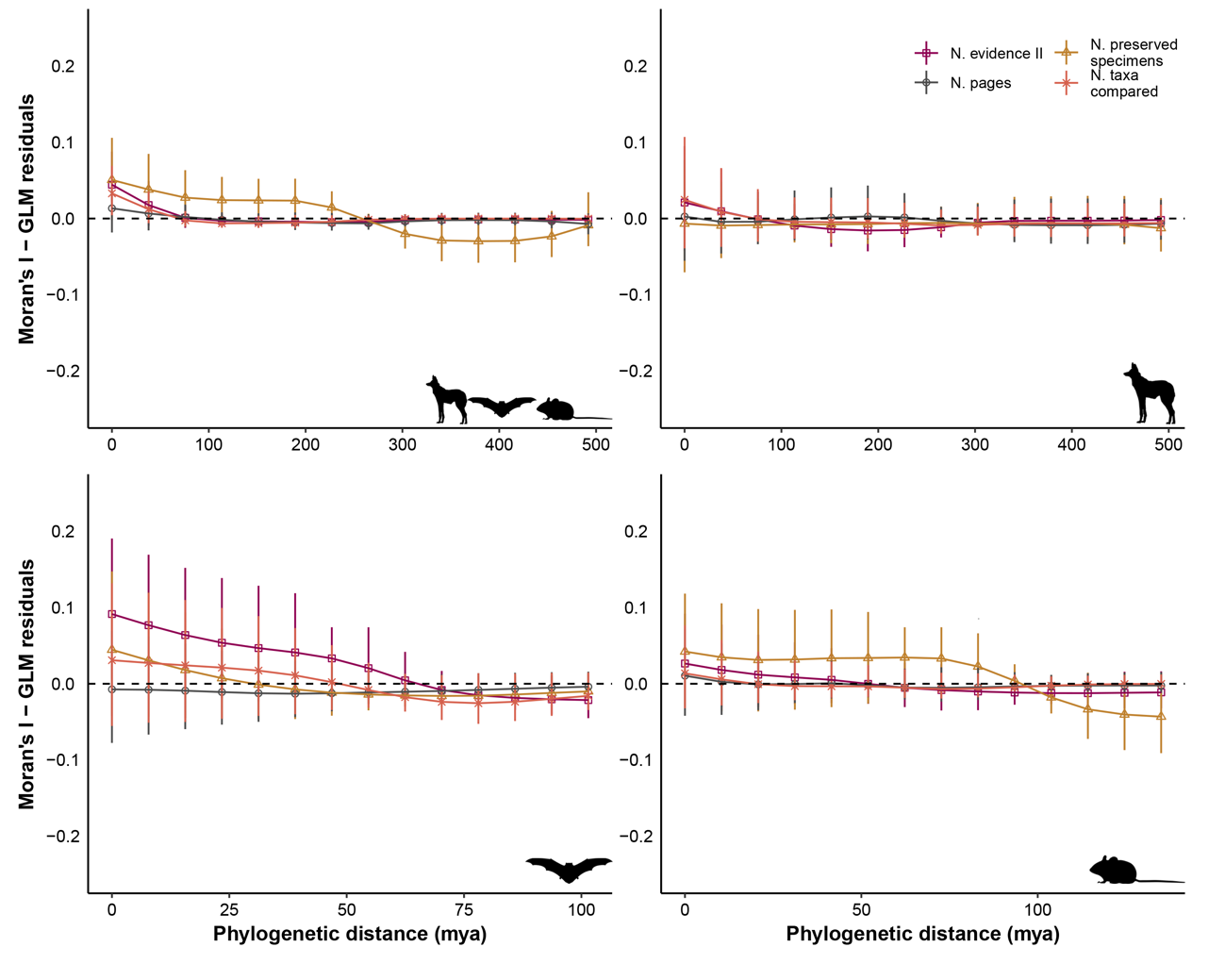

**Figure S5.** **Phylogenetic autocorrelation analysis of model residuals in (a) all mammals; (b) non-bats & non-rodents; (c) bats; and (d) rodents**. Moran’s *I* correlograms, which varies from -1 to +1, computed across 14 distance classes revealed maximum phylogenetic autocorrelations <0.2 (dashed horizontal line indicates null expectation), suggesting our regression models adequately accounted for phylogenetic structure. Lines represent 95% confidence intervals.

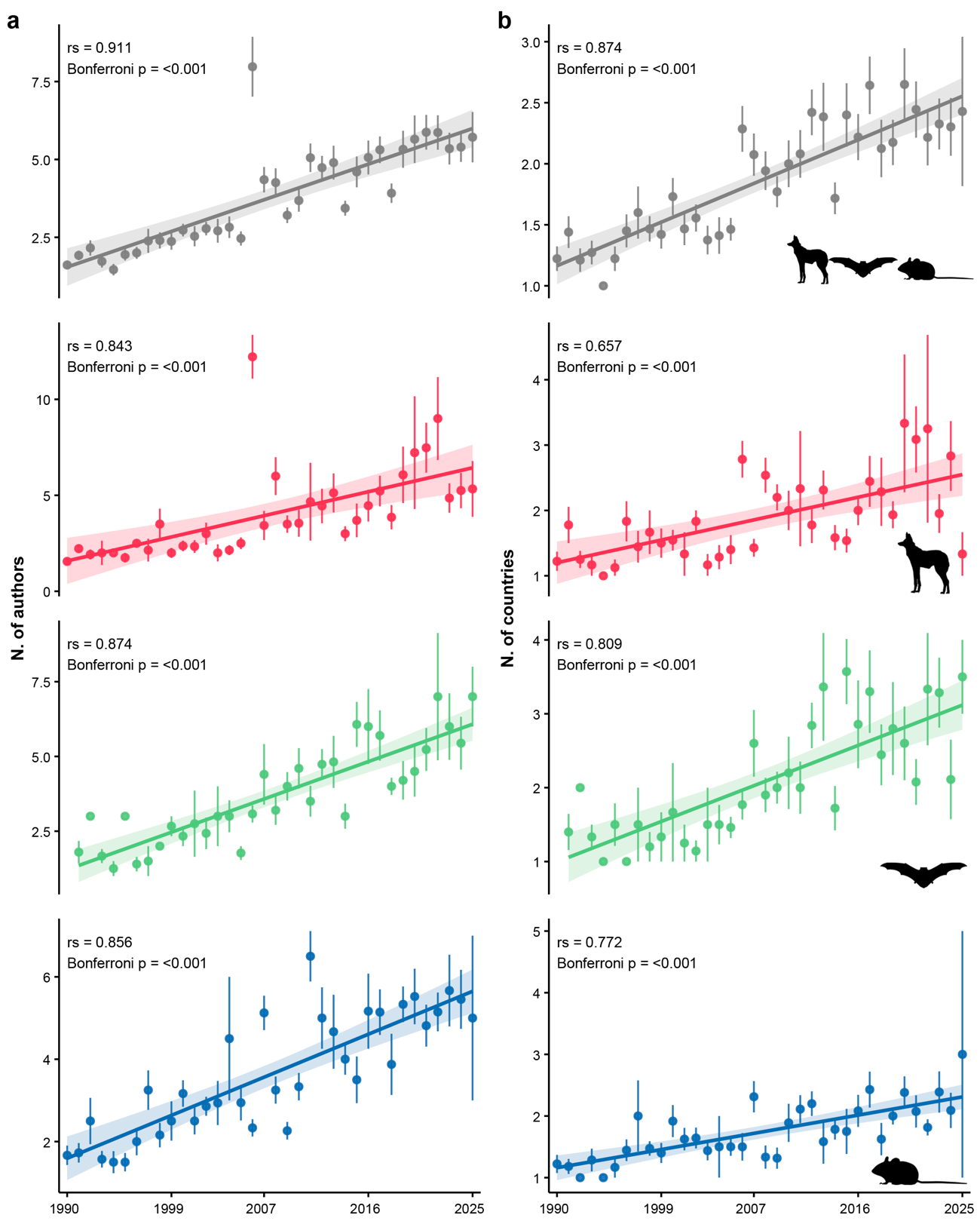

**Figure S6.** **Temporal trends in the number of authors and number of countries in mammal species descriptions from 1990–2025**. Panels show (a) all mammal species; (b) non-bats & non-rodents; (c) bats; and (d) rodents. Across all groups, there was an increase over time in the number of authors and countries of species descriptions.

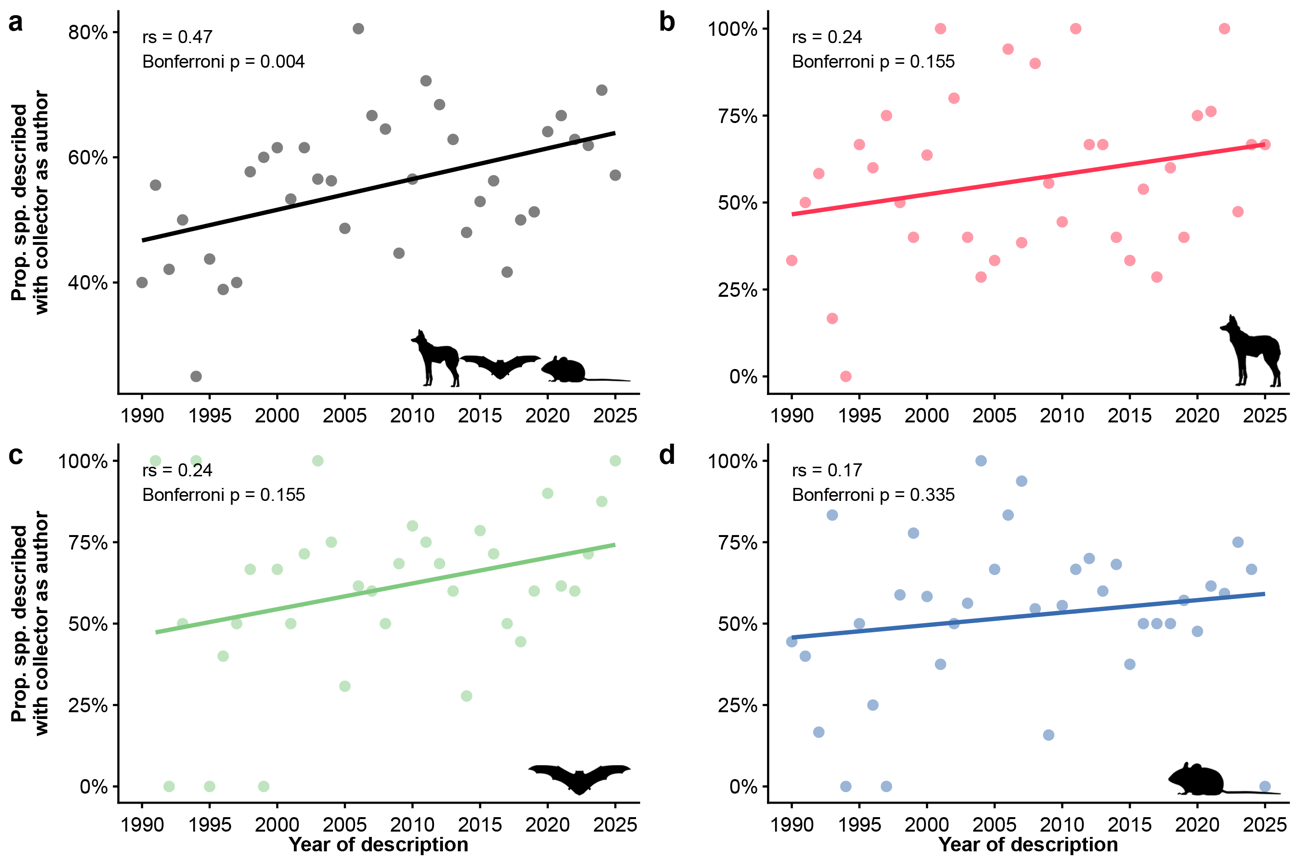

**Figure S7. Temporal trends in the proportion of mammal species whose holotype collector authored the formal species description**. Panels show (a) all mammal species; (b) non-bats & non-rodents; (c) bats; and (d) rodents. When we consider all mammals, there is an increase in the inclusion of collectors as authors of descriptions. In the other groups separately, there is only a positive tendency.
